# A discrete event simulation to evaluate the cost effectiveness of germline *BRCA1* and *BRCA2* testing in UK women with ovarian cancer

**DOI:** 10.1101/060418

**Authors:** Anthony Eccleston, Anthony Bentley, Matthew Dyer, Ann Strydom, Wim Vereecken, Angela George, Nazneen Rahman

## Abstract

**Objectives:** The objective of this study was to evaluate the long-term cost-effectiveness of germline *BRCA1* and *BRCA2* (collectively termed ‘BRCA’) testing in women with epithelial ovarian cancer, and testing for the relevant mutation in first and second degree relatives of BRCA mutation-positive individuals, compared with no testing. Female BRCA mutation-positive relatives of ovarian cancer patients could undergo risk-reducing mastectomy and/or bilateral salpingo-oophorectomy.

**Methods:** A discrete event simulation model was developed that included the risks of breast and ovarian cancer, the costs, utilities and effects of risk-reducing surgery on cancer rates, and the costs, utilities and mortality rates associated with cancer.

**Results:** BRCA testing all women with epithelial ovarian cancer each year is cost-effective at a UK willingness-to-pay threshold of £20,000/QALY compared with no testing, with an ICER of £4,339/QALY. The result was primarily driven by fewer cases of breast (142) and ovarian (141) cancer and associated reductions in mortality (77 fewer deaths) in relatives over the subsequent 50 years. Sensitivity analyses showed that the results were robust to variations in the input parameters. Probabilistic sensitivity analysis showed that the probability of germline BRCA mutation testing being cost-effective at a threshold of £20,000/QALY was 99.9%.

**Conclusions:** Implementing germline BRCA testing in all ovarian cancer patients would be cost-effective in the UK. The consequent reduction of future cases of breast and ovarian cancer in relatives of mutation-positive individuals would ease the burden of cancer treatments in subsequent years and result in significantly better outcomes and reduced mortality rates for these individuals.

## Introduction

Approximately 7,000 new cases of ovarian cancer are diagnosed in the UK every year [1–4], of which 13-16% are caused by a germline mutation in either the *BRCA1* or *BRCA2* (collectively termed ‘BRCA’) gene [5–9]. Knowing a patient’s BRCA mutation status is becoming increasingly important for optimal ovarian cancer management, providing information about response to chemotherapy, suitability for targeted agents such as poly adenosine diphosphate ribose polymerase (PARP) inhibitors, future cancer surveillance requirements and overall prognosis [10–13].

Women with a germline BRCA mutation have a 10-50% lifetime risk of ovarian cancer [14, 15] and a 40-85% lifetime risk of breast cancer [14, 15]. Because of this, relatives of BRCA mutation-positive individuals often undertake testing to find out if they have inherited the family mutation. This knowledge is used to decide whether to have enhanced cancer surveillance and/or risk-reducing surgery (RRS). If they choose to have RRS, bilateral salpingo-oophorectomy (RRBSO) and/or mastectomy (RRM) can be undertaken. Equally importantly, relatives that have not inherited the BRCA mutation can be spared these interventions.

Access to BRCA testing for ovarian cancer patients across the UK and Europe has been highly variable, with many centres using complex criteria to determine which patients should be offered testing. Historically, eligibility was primarily determined by family history of breast and/or ovarian cancer [16–18]. However, patients with a germline BRCA mutation do not always have a relevant family history of breast/ovarian cancer [5–7, 19], and therefore using these criteria to determine testing eligibility is sub-optimal.

The objective of this study was to determine the cost-effectiveness of providing germline BRCA mutation testing to all women with epithelial ovarian cancer in the UK, and the subsequent testing and management of their relatives that have a mutation. Of note, we have only considered germline BRCA mutations. The small proportion of ovarian cancer due to somatic BRCA mutations is not considered here; such mutations are not heritable and therefore do not have implications for relatives.

## Methods

### Model overview

A discrete event simulation model with annual cycles was developed in Microsoft Excel^®^. In the model, a simulated cohort of adult patients with ovarian cancer (index population) and their cancer-free family members were passed through various health states, including no cancer (family members only, with different risks of developing cancer depending on whether they choose RRS), ovarian cancer, breast cancer (family members only), and both ovarian and breast cancer. The model outputs were costs and quality-adjusted life years (QALYs), which were calculated for each individual and aggregated to provide an incremental cost-effectiveness ratio (ICER). The model also calculated the number of new cancer cases prevented and the number of lives saved. The flow of individuals through the model was based on defined characteristics, with the path determined by calculated time-to-events, or annual risks where time-to-event could not be calculated. The model adopted a 50-year time horizon, a UK health service perspective was used, and discount rates of 3.5% were applied to costs and outcomes, in accordance with UK health technology assessment guidelines [20].

The simulated index population consisted of 7,284 patients eligible for BRCA testing, which corresponds to the incidence of ovarian cancer in the UK in 2013 [4]. This population was included in two scenarios, BRCA testing or no BRCA testing, for the testing and non-testing arms.

Patients with a BRCA mutation entered the model (with mutation status known by testing or unknown in the non-testing arm). Patients who underwent BRCA testing but did not have a BRCA mutation did not enter the model, since there will be no difference in costs and outcomes between the testing and non-testing arms; however, the cost of testing these patients was included. Based on published data, 13% of patients were assumed to have a BRCA mutation, 60% of which were assumed to have a *BRCA1* mutation and 40% a *BRCA2* mutation [5–9]. Sensitivity analyses were included to vary this rate between 10% and 16%. If patients in the testing arm had a BRCA mutation, their simulated first-degree relatives were tested. If the relative had a BRCA mutation, simulated second-degree relatives were also tested. The age of simulated relatives upon model entry was calculated in relation to the age of the index case, and those aged <25 years were tested when they reached 25.

The model schematic is shown in Figure 1. An age of all-cause mortality was estimated for each individual using UK national life tables [21], and an annual age-adjusted risk of death was estimated for individuals with cancer, using published 5-year survival rates [22, 23]. Each year the model then determined whether individuals with cancer died from their cancer, until they reached their age of all-cause mortality.

RRS uptake was estimated using empirical data from The Royal Marsden hospital of 858 women, 458 with a *BRCA1* and 400 with *BRCA2* mutation. In *BRCA1* mutation carriers, the uptake of RRBSO was 88% and of RRM was 34%. In *BRCA2* mutation carriers, the uptake of RRBSO was 87% and of RRM was 25%. The uptake of RRBSO is slightly higher and the uptake of RRM slightly lower than published data [24, 25], therefore different rates of RRS uptake were included in sensitivity analyses. The age at which RRBSO occurred was assumed to be 40 years in *BRCA1* mutation-positive individuals and 45 years in *BRCA2* mutation-positive individuals, or on model entry for individuals above these ages. The age of RRM was assumed to be 40 years or on model entry for older individuals. The surgery cost and its impact on health-related quality of life (measured by a one-off disutility) were applied in the year that surgery took place. A hazard ratio (HR) was applied to the risk of cancer to reflect the lower risk after undergoing RRS.

When an individual developed cancer, treatment costs commenced and a risk of developing secondary cancer (breast/ovarian) was assigned. If secondary cancer developed, a new probability of age-adjusted cancer-related mortality was assigned.

**Figure 1:**
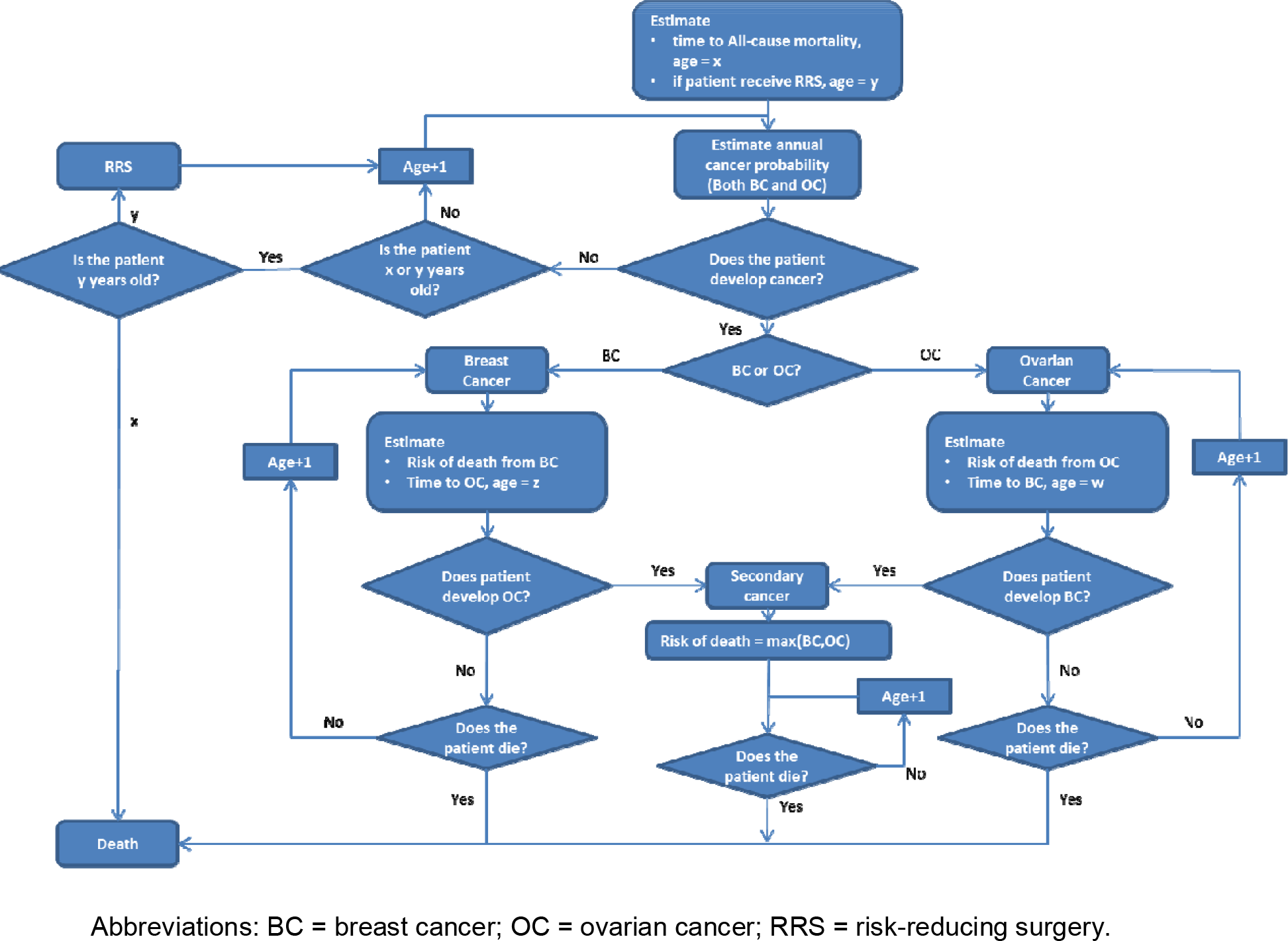
Model Schematic

### Data sources

The majority of data used in the model were UK-specific. Population data used to generate the model cohort are provided in (Table 1. Published UK data were used to estimate the mean number of siblings and children, as well as the mean age relative to the index case [21].

Cancer risk varied by age and BRCA mutation status (Table 2. A structured literature search was performed to identify the reduction in risk of breast cancer following RRM or RRBSO and the reduction in risk of ovarian cancer following RRBSO. There were eight relevant references [29–36], the data from which were used in a fixed effects meta-analysis to calculate the final HRs used in the model (Table 2. A fixed effects method was used rather than a random effects method due to low heterogeneity between studies. Only one publication [29] evaluated the risk reduction of breast cancer following both RRM and RRBSO. No evidence was identified to show that RRM affects the risk of ovarian cancer; therefore for patients undergoing both RRM and RRBSO the risk reduction of ovarian cancer following RRBSO was used.

The cancer-related mortality for both breast and ovarian cancer was estimated using 5-year net survival data reported by Cancer Research UK [22, 23], as shown in Table 2.

**Table 1:** Parameters for generating model cohort

| Index population | Data inputs |  | Number of patients |  | Reference |
| --- | --- | --- | --- | --- | --- |
| Number of cases | 7,284 |  | 7,284 |  | ONS [1] |
| Mean age, years (SD<sup>†</sup>) | 50 (5) |  | – |  | Domchek 2013 [26] |
| BRCA mutation | 13% |  | 964 |  | Risch et al [8]<br>Zhang et al[9]<br>Alsop et al [5]<br>George et al [6]<br>Pal et al [7] |
| Proportion with *BRCA1* mutation | 60% |  | 583 |  |  |
| Proportion with *BRCA2* mutation | 40% |  | 381 |  |  |
| First-degree relatives | Mother | Father | Siblings | Children | Reference |
| Mean number (SD<sup>†</sup>) | 1 | 1 | 0.91 (0.5) | 1.91 (0.5) | ONS [27] |
| Mean age relative to index case (SD<sup>†</sup>) | 30 (5) | 32 (5) | 0 (5) | –30 (%) | ONS [27] |
| Sex, probability female | 100% | 0% | 50.78% | 50.78% | ONS [28] |
| Probability BRCA mutation | 50% | 50% | 50% | 50% | – |
| Second-degree relatives | Grand-parents | Uncles/aunts | Nieces/nephews | Grand-children | Reference |
| Mean number (SD<sup>†</sup>) | 4 | 1.8 (0.5) | 1.7 (0.5) | 3.6 (0.5) | Calculation based on ONS [27] |
| Mean age relative to first-degree relative (SD<sup>†</sup>) | 30 (5) | 0 (5) | –30 (5) | –30 (5) | ONS [27] |
| Sex, probability female | – | 50.78% | 50.78% | 50.78% | ONS [28] |
| Probability BRCA mutation | 25% | 25% | 25% | 25% | – |

### Costs

Costs were included for BRCA testing, genetic counselling, cancer surveillance, RRS, hormone replacement therapy (HRT), cancer treatment, and palliative care (Table 3). HRT was included for individuals undergoing RRBSO without a history of breast cancer until the age of 52 years, as recommended by NICE guidelines [17]. For genetic counselling, one post-test session for index patients with a BRCA mutation, one pre-test genetic session for all relatives, and one additional post-test session for relatives found to have a BRCA mutation were included. This is in accordance with the mainstream model of genetic testing used at The Royal Marsden [6]. In sensitivity analyses, relatives received two pre-test counselling sessions as recommended by NICE [17].

Costs for BRCA testing, genetic counselling, and RRS were applied in the cycle in which they occurred, whereas costs for HRT and surveillance (MRI and mammography) were applied annually; HRT costs were applied after RRBSO until the age of 52 or the development of breast cancer, and MRI and mammography costs were applied after BRCA testing in BRCA mutation-positive patients until either breast or ovarian cancer developed.

Cancer treatment costs were derived from a micro-costing exercise conducted in 2013 for the NICE familial breast cancer guideline [37]. Given the short life expectancy of those developing ovarian cancer and the high likelihood of repeat treatment, costs of treating ovarian cancer were applied annually. The survival rate for breast cancer is much greater, and therefore it was assumed that all treatment costs for breast cancer were applied for one year during the cycle where diagnosis occurred; however, it is acknowledged that breast cancer treatment may last longer. It was assumed that individuals who received a mastectomy before breast cancer diagnosis did not require surgery as part of their treatment; however, patients with ovarian cancer after RRBSO were assumed to require additional de-bulking surgery. Palliative care costs were applied in the cycle that the patient died.

**Table 2:** Cancer risks, risk reduction following RRS and 5-year cancer survival rates

| Approximate 5-year risk of breast cancer by age |  |  | Approximate 5-year risk of ovarian cancer by age |  |  |
| --- | --- | --- | --- | --- | --- |
| Age range, years | BRCA1 | BRCA2 | Age range, years | BRCA1 | BRCA2 |
| 20–25 | 5% | ~1% | 30–39 | 5% | 5% |
| 26–30 | 5% | 2% | 40–44 | 10% | 5% |
| 31–35 | 5% | 5% | 45–49 | 10% | 10% |
| 36–40 | 10% | 2% | 50–54 | 15% | 10% |
| 41–45 | 10% | 10% | 55–59 | 10% | 10% |
| 46–50 | 15% | 10% | 60–64 | 10% | 5% |
| 51–55 | 15% | 10% | 65–69 | 10% | 5% |
| 56–60 | 10% | 10% | 70–79 | 10% | 5% |
| 61–65 | 10% | 15% |  |  |  |
| 66–70 | 10% | 15% |  |  |  |
| Hazard ratios of RRS from meta-analysis - BRCA1 |  |  | Hazard ratios of RRS from meta-analysis - BRCA2 |  |  |
| RRS | Breast cancer HR (95% CI) | Ovarian cancer HR (95% CI) | RRS | Breast cancer HR (95% CI) | Ovarian cancer HR (95% CI) |
| RRM | 0.10 (0.03–0.31) | 1.00 | RRM | 0.09 (0.03–0.31) | 1.00 |
| RRBSO | 0.51 (0.39–0.66) | 0.16 (0.09–0.26) | RRBSO | 0.39 (0.29–0.54) | 0.12 (0.06–0.23) |
| RRM + RRBSO | 0.05 (0.01–0.22) | 0.16 (0.09–0.26) | RRM and RRBSO | 0.05 (0.01–0.22) | 0.12 (0.06–0.23) |
| 5-year net survival rate for breast cancer |  |  | 5-year net survival for ovarian cancer |  |  |
| Age range, years | % age Survival |  | Age range, years | % age Survival |  |
| 15–39 | 84.9 |  | 15–39 | 87.4 |  |
| 40–49 | 90.0 |  | 40–49 | 74.0 |  |
| 50–59 | 91.2 |  | 50–59 | 59.6 |  |
| 60–69 | 92.4 |  | 60–69 | 43.0 |  |
| 70–79 | 83.0 |  | 70–79 | 35.7 |  |
| 80–99 | 70.3 |  | 80–99 | 20.4 |  |

Abbreviations: CI = confidence interval; HR = hazard ratio; RRBSO = risk-reducing bilateral salpingo-oophorectomy; RRM = risk-reducing mastectomy; RRS = risk-reducing surgery.

**Table 3:** Costs

| BRCA testing, RRS and surveillance |  | Cost | Reference |  |  |
| --- | --- | --- | --- | --- | --- |
| BRCA testing | Index case (full genes) | £306 | Royal Marsden [38] |  |  |
|  | Family members (specific mutation only) | £108 | Royal Marsden [38] |  |  |
|  | Genetic counselling, per 2-hour session | £126 | NICE CG164 [37]: Based on rate per hour of patient contact for band 7 counsellor in primary medical care |  |  |
| RRS | Mastectomy including reconstructive surgery | £9,219 | NHS reference costs 2014–15 [39]: Weighted average of HRG codes JA27Z and JA28Z |  |  |
|  | BSO | £2,976 | NHS reference costs 2014–15 MA08A–MA08B [39] |  |  |
|  | HRT, per year | £120.95 | BNF 69 2015 [40] and HSCIC Prescription Cost Analysis 2014 [41]: Weighted average of Klioavance, Evorel Conti and Evorel Sequi |  |  |
| Surveillance |  |  |  |  |  |
| MRI, per year |  | £191 | NHS reference costs 2014–15 [39]: HRG code RA05Z |  |  |
| Mammography, per year |  | £55 | NICE CG144 costing report for venous thromboembolic diseases [42]; uplifted using PSSRU unit costs of health and social care 2014 [43] |  |  |
| Treatment |  | Unit cost | Dose/units | Total cost | Reference |
| Breast cancer | Breast surgery | £3,186 | 1 | £3,816 | NHS reference costs 2014–15 [39]: Weighted average of HRG codes JA28A–C, JA39Z–JA41Z |
|  | Adjuvant radiotherapy | £132 | 15 | £1,978 | NHS reference costs 2014–15 [39]: HRG code SC23Z |
|  | Chemotherapy delivery: first attendance | £389 | 1 | £389 | NHS reference costs 2014–15 [39]: HRG code SB14Z |
|  | Chemotherapy delivery: subsequent attendance | £326 | 5 | £1,632 | NHS reference costs 2014–15 [39]: HRG code SB15Z |
|  | Chemotherapy drugs (fluorouracil, epirubicin, cyclophosphamide) | £205 | 6 | £1,230 | BNF 69 2015 [40] |
|  | Neulasta<sup>†</sup> | £686 | 6 | £4,118 | BNF 69 2015 [40] |
|  | Dexamethasone<sup>§</sup> | £0.78 | 16 mg OD for 2 days | £12 | BNF 69 2015 [40] |
|  | Anastrozole<sup>¶</sup> | £0.07 | 1 mg OD for 5 years | Variable<sup>‡</sup> | BNF 69 2015 [40] |
|  | Total with surgery |  |  | £13,189 | – |
|  | Total without surgery |  |  | £9,373 | – |
| Ovarian cancer | De-bulking surgery | £5,613 | 1 | £5,613 | NHS reference costs 2014–15 MA26A–MA26C [39] |
|  | Chemotherapy delivery: first attendance | £389 | 1 | £389 | NHS reference costs 2014–15 [39]: HRG code SB14Z |
|  | Chemotherapy delivery: subsequent attendance | £326 | 5 | £1,632 | NHS reference costs 2014–15 [39]: HRG code SB15Z |
|  | Chemotherapy drugs (33% carboplatin, 67% carboplatin + paclitaxel) | £568 | 6 | £3,408 | BNF 69 2015 [40] |
|  | Neulasta<sup>†</sup> | £668 | 6 | £4,118 | BNF 69 2015 [40] |
|  | Dexamethasone<sup>§</sup> | £0.78 | 16 mg OD for 2 days | £12 | BNF 69 2015 [40] |
|  | Total with surgery |  |  | £15,185 | – |
|  | Total without surgery |  |  | £9,572 | – |
| Palliative care |  |  |  |  |  |
| Condition |  | Cost | Reference |  |  |
| Breast cancer |  | £3,702 | UK study of treatment patterns and resource costs for specific advanced cancer patients [44]; uplifted to 2013–14 costs from PSSRU [43] |  |  |
| Ovarian cancer |  | £7,143 |  |  |  |
| All-cause mortality |  | £103 | NHS reference costs 2014–15 [39]: HRG code SD03A |  |  |

BNF = British National Formulary; BSO = bilateral salpingo-oophorectomy; HRG = Healthcare Resource Group; HR = hormone replacement therapy; HSCIC = Health and Social Care Information Centre; MRI = magnetic resonance imaging; NICE = National Institute for Health and Care Excellence; OD = one daily; PSSRU = Personal Social Services Research Unit; RRS = risk-reducing surgery. <sup>†</sup>Used to treat neutropenia to reduce the risk of infection. <sup>§</sup>Used to treat inflammation, relieve sickness and boost appetite. <sup>‡</sup>Used to inhibit the synthesis of oestrogen as adjuvant treatment in oestrogen-receptor positive breast cancer. <sup>‡</sup>The total cost of anastrozole varies between patients, since some patients may die within the 5 years specified to receive this medication.

**Table 4:** Utility Values

| Weighted health state index by age group in females |  |  |
| --- | --- | --- |
| Age, years | Utility, mean (SD) |  |
| <25 | 0.94 (0.12) |  |
| 25–34 | 0.93 (0.15) |  |
| 35–44 | 0.91 (0.15) |  |
| 45–54 | 0.85 (0.26) |  |
| 55–64 | 0.81 (0.26) |  |
| 65–74 | 0.78 (0.25) |  |
| ≥75 | 0.71 (0.27) |  |
| Cancer-related utilities |  |  |
| Time from diagnosis | Ovarian cancer | Breast cancer |
| Year 1 | 0.50 | 0.71 |
| Year 2 | 0.65 | 0.72 |
| Year 3 | 0.67 | 0.73 |
| Year 4 | 0.69 | 0.74 |
| Year 5 | 0.70 | 0.76 |
| Year 6+ | 0.72 | 0.77 |
| Health state utilities |  |  |
| Health states | Utility values in controls, mean (SD) | Utility values in BRCA mutation carriers, mean (SD) |
| Perfect health | 1.00 | 1.00 |
| RRM | 0.88 (0.17) | 0.88 (0.22) |
| RRBSO | 0.90 (0.14) | 0.95 (0.10) |
| Both RRSs | 0.79 (0.21) | 0.84 (0.23) |
| HRT | 1.00 | 1.00 |
| Healthy with a known BRCA mutation (sensitivity analysis only) | 0.87 (0.16) | 0.92 (0.15) |
| Death | 0.00 | 0.00 |

### Health state utilities

Age-related utilities for females [45] were used in the model to ensure that the QALY gain associated with BRCA testing was not overestimated (Table 4).

The NICE clinical guideline 164 cost-effectiveness evidence review [46] provided utilities for both ovarian and breast cancer following diagnosis. These disease-specific utilities were combined with the age-related utilities multiplicatively as advised by the NICE Decision Support Unit [47], and the impact on quality of life was assumed to decrease each year after diagnosis until Year 6, after which it remained constant. If a patient was diagnosed with both cancers, the utility values were also applied multiplicatively.

Utility values for the other health states and treatments in the model were derived from a time trade-off study in BRCA mutation-positive individuals [48]. This study reported that RRS was associated with a short-term detrimental impact on HRQoL and, consistent with UK clinical opinion, these utility values were assumed to apply only in the cycle in which RRS occurred. The base case analysis applied no disutility for having a BRCA mutation, which is consistent with The Royal Marsden experience and other published studies [49, 50]. However, a disutility has been reported in at least one study [48] and thus was included in sensitivity analyses.

### Sensitivity analyses

Parameter uncertainty around key model inputs was tested using sensitivity analyses, where parameters were independently varied over a plausible range determined by either the 95% confidence interval or by clinical expert opinion; where no estimates were available a range of ±25% was used.

Joint parameter uncertainty was also explored through probabilistic sensitivity analysis (PSA), where all parameters were assigned distributions and varied jointly.

### Model assumptions

There were a number of assumptions made during the development of the model:

- The sensitivity and specificity of full BRCA gene and specific mutation testing was 98%. This corresponds with Royal Marsden empirical data and published literature [51, 52].
- Relatives with a BRCA mutation had the same BRCA mutation as the index case.
- Relatives considered in the model had no prior ovarian or breast cancer and had not undergone RRS.
- The 5-year and 10-year risks for breast cancer and ovarian cancer, respectively, were constant over the 5 or 10 years. This is a simplifying assumption arising from the 5-year and 10-year risk data used in the model for breast cancer and ovarian cancer.
- All RRMs were bilateral. This is a simplifying assumption arising because the HRs obtained from the literature were reported for patients receiving bilateral mastectomy.
- Patients did not develop both breast and ovarian cancer in the same year. This is a simplifying assumption supported by Royal Marsden data. Although clinically possible, it is extremely unusual.
- The index population did not receive RRM. This is a simplifying assumption, since only a small number of ovarian cancer patients with BRCA mutations have RRM.
- The costs and outcomes for patients without a BRCA mutation were equal between the testing and non-testing arms, because the risks of developing breast and/or ovarian cancer were the same for these patients in both arms. This means that the model only considers the incremental difference between testing and no testing in BRCA mutation-positive individuals (although the cost of testing individuals without a BRCA mutation was included).
- The population was not dynamic; therefore the model did not consider relatives born after the index case was tested. This was a simplifying assumption because a dynamic population would have been impractically complex to model. However, the approach taken allowed the results for testing an incident population from a single year to be assessed; the benefits of testing would be seen over the lifetime of these patients regardless of whether the testing scheme continued for longer than one year.
- The model was not a typical oncology cost-utility model, and did not specifically consider treatments received or cancer severity.

## Results

### UK base case

There were 7,284 index cases run through the model, resulting in 3,768 first-degree and 935 second-degree family members eligible for testing. In total, BRCA testing identified 1,314 patients with a *BRCA1* mutation and 886 with a *BRCA2* mutation (Table 5).

The total discounted cost of BRCA testing (£9.6m) was partially offset by a reduction in cancer treatment and palliative care costs, leading to an incremental discounted cost of £3.0m. Over the 50-year time horizon, there were an additional 706 discounted QALYs associated with BRCA testing compared with no testing, resulting in an ICER of £4,339/QALY, which is well below the UK threshold of £20,000/QALY.

An important consequence of implementing BRCA testing in ovarian cancer patients is the reduction in cancer and deaths in their relatives. If all women diagnosed with ovarian cancer were tested in one year, this analysis has calculated that there would be 77 fewer deaths, 141 fewer new cases of ovarian cancer, and 142 fewer new cases of breast cancer in relatives over 50 years.

It is also important to note that BRCA testing provided no benefit to index cases in this model, since they entered the model with ovarian cancer and did not undergo RRM. Therefore, benefits were only seen in the first and second degree relatives who have BRCA testing before developing breast and/or ovarian cancer and have the option of RRS. This can be seen as a conservative assumption, since it is possible that patients in the index population may undergo RRM.

**Table 5:**
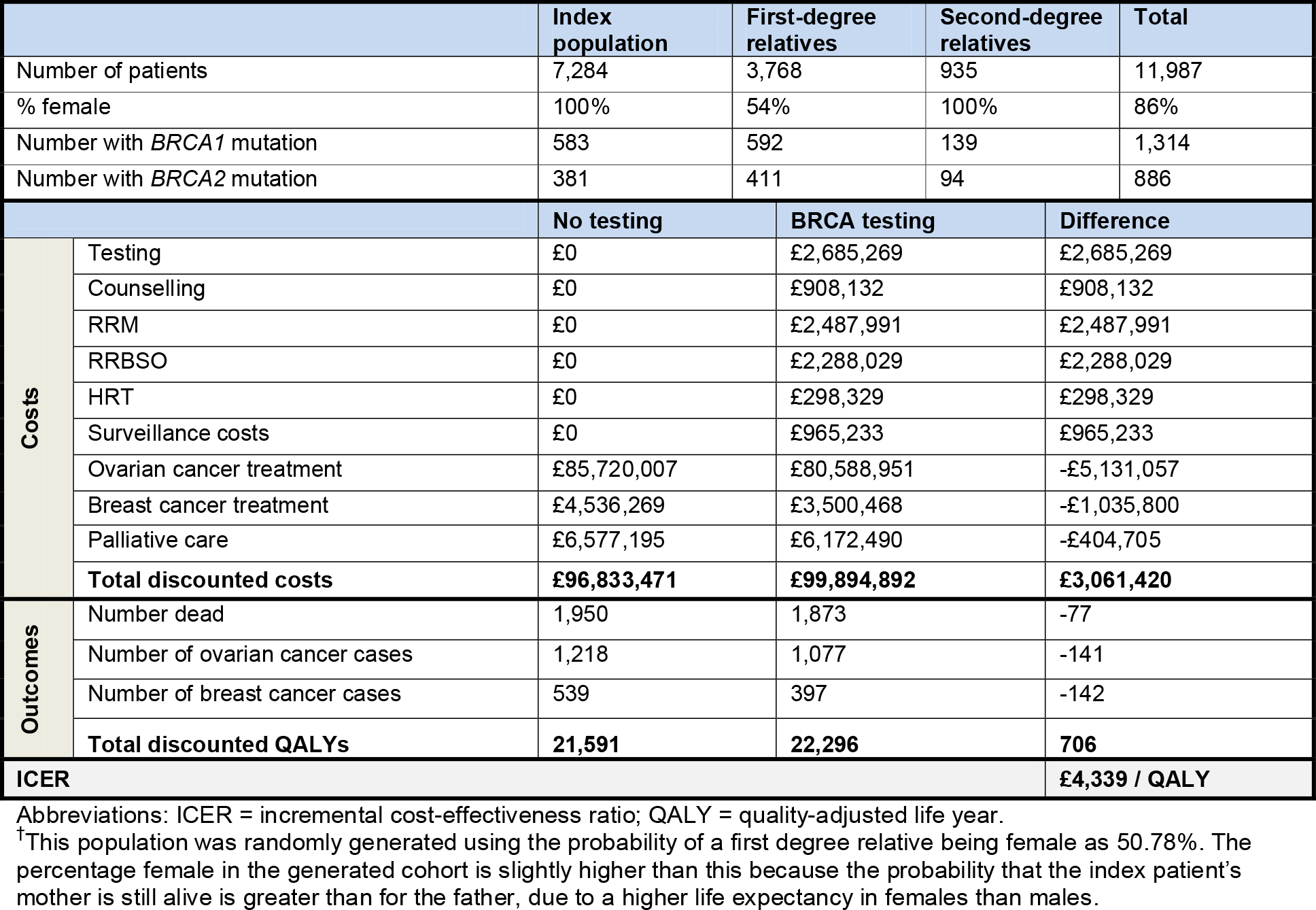
Cost-effectiveness results

### Sensitivity analyses

The results from the one-way sensitivity analyses did not differ substantially from the base case, and all results were below the UK cost-effectiveness threshold of £20,000/QALY.

Changing the probability of having a BRCA mutation to 10% and 16% (base case 13%) had little effect on the ICER (£5,975 and £6,373/QALY, respectively).

The RRBSO uptake rate in some published data [24, 25] is lower than Royal Marsden data, and lowering the RRBSO uptake rate to 75% increased the ICER to £6,139/QALY. Conversely, RRM uptake in published data is higher than Royal Marsden data [24, 25]. Increasing the RRM uptake rate to 50% resulted in a slightly higher ICER (£5,353/QALY) than the base case, because the higher costs of treatment were not offset by survival gains, due to high breast cancer survival in patients who do not undergo RRM.

Increasing the mean age of the index population to 60 years lowered the ICER to £3,811/QALY. This was due to the generation of more grandchildren, and so there were more relatives receiving RRS and therefore more quality-adjusted life years were accrued. Conversely, decreasing the mean age to 40 years increased the ICER to £4,481/QALY.

Using the 95% confidence intervals for the HRs for risk reductions in developing cancer following RRBSO resulted in ICERs that were similar to the base case (£3,480 and £6,449/QALY), while using the 95% confidence intervals for RRM did not change the ICER (when accounting for rounding). This is because the confidence interval ranges for RRM are very small and therefore have a very small effect on the ICER. Increasing the survival rates by 25% resulted in a higher ICER of £4,442/QALY, while a decrease of 25% led to a lower ICER (£4,165/QALY). However, for ovarian cancer, 25% higher survival rates led to a lower ICER (£3,458/QALY) and 25% lower survival resulted in a higher ICER (£5,399/QALY). This is because ovarian cancer costs are so high, and despite a greater QALY gain for BRCA testing with a lower survival rate, the cost saving associated with fewer ovarian cancer cases is lower.

Including two pre-test genetic counselling sessions for relatives of the index population, as per NICE guidelines [17], slightly increased the ICER to £5,094/QALY. When a disutility associated with BRCA testing of 0.87 was applied, this resulted in fewer QALYs gained (508) and a slightly higher ICER of £6,026/QALY.

Probabilistic sensitivity analysis (5,000 simulations of the cohort) showed that the expected ICER was £5,282/QALY (95% CI: £1,593, £11,764). All simulation results were in the north-east or south-east quadrant of the cost-effectiveness plane, meaning that BRCA testing was always more effective than no testing. Overall, the probability of BRCA testing being cost-effective using a £20,000/QALY threshold was 99.9%.

## Discussion

This study shows that implementing routine BRCA testing in women with ovarian cancer would be cost-effective in the UK compared with no testing. It would result in lower breast and ovarian cancer incidence rates, lower treatment costs, lower cancer-related mortality, and an overall higher quality of life. The lives saved and the fewer new cases of ovarian and breast cancer in relatives in the testing arm are particularly important results in driving implementation.

NICE and the Cancer Strategy Taskforce recommend that cancer patients at >10% risk of having a BRCA mutation should be offered testing [17, 53]. Several recent studies have shown that any woman with epithelial ovarian cancer is eligible by this criterion [5–9]. Many centres use family history of cancer to determine test eligibility, but this is much less effective in identifying women with BRCA mutations [54]. Some centres restrict testing to non-mucinous or high-grade serous ovarian cancer. However, only ~3% of ovarian cancers are mucinous [55], some of which are due to other cancer predisposition genes that are frequently concurrently tested with *BRCA1* and *BRCA2* [56]. Therefore, it is simplest to offer testing to all women with epithelial ovarian cancer. This would likely require some additional funding, although the increase in the number of tests will in part be offset by the substantial recent decrease in the cost of testing due to the use of new sequencing technologies [57]. Furthermore, as our results show, there will be longer-term cost and health benefits. While this analysis was to calculate the cost-effectiveness of BRCA testing versus no testing and therefore included all eligible patients, it is acknowledged that the uptake rate of BRCA testing may not be 100% in clinical practice.

Within a cost-effectiveness analysis there are multiple sources of uncertainty, e.g. parameter and structural, which must be accounted for to increase the confidence that can be placed in the model output. Structural uncertainty occurs when there is uncertainty around the functional form of the model and whether it adequately reflects the patient pathway; while parameter uncertainty occurs when the exact value of an input is unknown. Standard modelling approaches, clinician validation and sensitivity analysis have all been used to reduce and understand the uncertainty and bias within this model.

There were a number of limitations associated with the model and the data inputs used. This was not a typical oncology cost-utility model that tracks overall survival and progression-free survival, and it therefore did not specifically consider the treatments received, and no variation in cancer severity has been modelled. However, the model used survival rates and average costs to reflect the impact of BRCA mutation testing and subsequent RRS, and including cancer staging and alternative treatments would have made the model impractically complex.

The simplified methodology of this model means that only the relatives of the index case benefit from BRCA testing. However, BRCA testing may benefit many patients with ovarian cancer because mutation-positive individuals typically have enhanced breast surveillance and some elect to have RRM. Additionally, BRCA mutation-positive ovarian cancer patients are increasingly able to access targeted therapies such as PARP inhibitors. PARP inhibitor therapies have also been shown to have activity in breast cancer [58, 59] and in male patients with BRCA mutation-positive prostate cancer [60, 61]. In our model, male first-degree relatives were tested for the BRCA mutation to identify any second-degree female relatives for testing; however no benefit to them was taken into account. The knowledge of BRCA mutation may provide patients with breast, ovarian and prostate cancer access to targeted therapies that would not benefit patients without a BRCA mutation.

Another limitation is that no mortality or morbidity was considered for RRS; although this may bias the analysis in favour of testing, the rates of mortality and morbidity are generally low [31].

Patients with a BRCA mutation who choose not to receive RRS are still eligible for increased surveillance; NICE clinical guidance 164 for familial breast cancer recommends that BRCA mutation carriers aged 30-49 years should undergo annual magnetic resonance imaging (MRI) surveillance, and those aged >40 years should have annual mammograms [17]. It is important to note that it was not possible to capture the benefit of increased surveillance in terms of earlier diagnosis of cancer, because the analysis did not specifically consider patients at different stages of their disease; however the extra surveillance costs have been included and therefore the results can be considered conservative.

The NHS also recommend screening mammograms every 3 years in all women aged 50-70 years [17]. This was not included in the non-testing arm of the model for women who had a BRCA mutation. Again, this can be seen as a conservative assumption, since including the costs of screening patients who were unaware of the mutation would increase the costs in the non-testing arm, and therefore reduce the incremental costs between the two arms and reduce the ICER, making BRCA testing even more cost-effective.

A previously published study by Kwon et al in 2009 [62] estimated the cost-effectiveness of BRCA mutation testing in women with ovarian cancer in the USA, and the downstream benefits for the first-degree relatives of patients with a BRCA mutation from the option of undergoing RRS. The study found that BRCA testing of women with ovarian cancer and a personal history of breast cancer, a family history of breast/ovarian cancer, or of Ashkenazi Jewish ancestry, was cost-effective by preventing future breast and ovarian cancers among first-degree relatives with an ICER of $32,018 per life year gained compared with no testing. This study cannot be directly compared with our results due to a number of differences, e.g. the analysis was based on US payer perspective, and BRCA testing was only performed on patients with a personal or family history of cancer or Ashkenazi Jewish ancestry. However, both studies concluded that BRCA mutation testing with the option of RRS in relatives of patients with a BRCA mutation was cost-effective compared with no testing.

## Conclusion

The base-case analysis results show that germline BRCA mutation testing in women with epithelial ovarian cancer is cost-effective at a UK threshold of £20,000/QALY compared with no testing, with an ICER of £4,339/QALY. If all ovarian cancer patients are tested in one year, there would be 141 fewer new cases of ovarian cancer, 142 fewer new cases of breast cancer, and 77 fewer deaths. These findings are robust to changes in the parameters, with all sensitivity analyses producing an ICER below £20,000/QALY, and the probability that BRCA testing is cost-effective at this threshold is 99.9%. Implementing BRCA testing for all women with ovarian cancer would require some reorganisation of testing services and may have some upfront resource implications; however, the reductions in the number of cases of both breast and ovarian cancer would ease the burden of cancer treatments in subsequent years and result in reduced mortality rates for these cancers.

